# Are Zinc oxide nanoparticles safe? A structural study on human serum albumin using *in vitro* and *in silico* methods

**DOI:** 10.1101/503797

**Authors:** Marziyeh Hassanian, Hassan Aryapour, Alireza Goudarzi, Masoud Bezi Javan

## Abstract

With due attention to adsorption of proteins on the nanoparticles surface and the formation of nanoparticle-protein corona, investigation of nanoparticles toxicity on the structure of proteins is important. Therefore, this work was done to evaluate toxicity of Zinc oxide nanoparticles (ZnO NPs) on the structure of human serum albumin (HSA) through *in vitro* and *in silico* studies. First, ZnO NPs were synthesized using hydrothermal method and their size and morphology were determined by SEM and TEM methods and then to study its toxicity on the HSA structure were used UV-Vis and fluorescence spectroscopy. Also, in order to investigate interaction mechanism of ZnO NP with HSA at the atomistic level was used molecular dynamics (md) simulation. The obtained images from SEM and TEM showed that ZnO NPs were nanosheet with size of less than 40 nm. The results of spectroscopic studies showed ZnO NPs lead to significant conformational changes in the protein’s absorption and emission spectra. Moreover, md results showed the minor structure changes in HSA due to interaction with ZnO NP during the 100 ns simulation and the formation of nanoparticle-protein corona complex that is mainly because of electrostatic interactions between charge groups of HSA and ZnO NP.

## 1. Introduction

Nanotechnology is the ability to study and use of particles in dimensions between 1-100 nm which commonly known as nanoparticles (NPs). NPs have unique mechanical and chemical properties arising from their high surface area and quantum effects. Therefore, NPs are used in a wide range of different applications and humans can be exposed to NPs via several routes such as inhalation, injection, oral ingestion and the dermal route ^[1]^. Upon the entry of NPs into the human body, they may interact with biomacromolecules and even biological metabolites due to their nano size and large surface to mass ratio. But, the adsorption of proteins on the surface of NPs and the formation of nanoparticle-protein complex which is known as the nanoparticle-protein corona (NP-PC), is unique ^[2]^. On the other hand, adsorption of proteins on the NPs surface may result in structural changes and unfolding of proteins and lead to toxicity ^[3]^. The previous studies showed that the binding of ZnO NPs led to the denaturation of the periplasmic domain of ToxR protein of *Vibrio cholerae* ^[4]^. Therefore, it is important to study the toxicity of NPs on the structure of proteins to design nanoparticle that do not have toxic effects on the structure of proteins and consequently, on living organisms. In this study, we synthesized ZnO NPs and studied their toxicity on human serum albumin (HSA). These nanoparticles are an important class of commercially applied material and used in diagnostics, therapeutics, drug delivery systems, electronics, cosmetics, personal care products, food additives and pigments and coatings due to their magnetic, catalytic, semiconducting, antimicrobial, ultraviolet protective and binding properties ^[5]^. Moreover, human serum albumin is the most abundant type of plasma protein (3.5 to 5.5 g/dL) with highest electrophoretic mobility, and it has a globular structure with a molecular weight 66.5 kDa, consists of 585 amino acids, 17 disulfide bonds, and only one free thiol group (Cys34) ^[6]^. HSA comprises three homologous domains (I, II, and III) that assemble to form a heart-shape molecule and each domain is divided into two subdomains A and B ^[7]^. The function of HSA is the maintenance of colloid osmotic pressure and transportation, distribution and metabolism of many endogenous and exogenous small molecules and ions ^[8, 9]^. Thus, due to the increasing use of ZnO NPs in various products and also the biomedical importance of HSA in human health, we investigated the toxicity of ZnO NPs on the structure of this protein through *in vitro* and *in silico* studies.

## 2. Materials and Methods

### 2.1. Materials

Zinc acetate dihydrates [(Zn (CH_3_COO)_2_.2H_2_O)], ti-sodium citrate and sodium hydroxide were purchased from Merck Company and Human serum albumin (96%) and ANS (1-anilino-8-naphthalene sulfonic acid) were purchased from Sigma-Aldrich.

### 2.2. Synthesis of ZnO NPs

ZnO NPs were synthesized using hydrothermal method as follows ^[10]^. First, 60 ml of 1 M Zinc acetate and 30 mL of 0.5 M citrate were mixed in a baker, then 50 mL distilled water and 60 ml of 1 M sodium hydroxide were added dropwise to the mixed solution along with sonication. The obtained white solution was incubated at room temperature for 24 h, and then the resulting pellet was washed with distilled water several times, and lyophilized to give nanopowder. The size and morphology of ZnO NPs were determined by SEM (Hitachi S-4160) and TEM (Philips CM 120).

### 2.3. Spectroscopy procedures

All experiments were performed in 0.1 mM sodium phosphate buffer (pH 7.4), and different concentration of ZnO NPs (0, 0.1, 0.2, 0.3, 0.4 and 0.5 mg/mL) were incubated with a constant concentration of HSA (5 μM) for 1 h at room temperature. The UV-Vis absorption spectra were recorded with double-beam Shimadzu UV-1800 UV-Vis spectrophotometer. After incubation of HSA with different concentrations of ZnO NPs, the absorption spectra were taken in the wavelength range of 200-500 nm. Time kinetics study was done using a BioTek ELx808IU microplate reader at pH 7.4 to determine the time depended absorption of HSA interaction (5 μM) with ZnO NP (0.5 mg/mL) under temperature 37 °C for 120 min.

Also, fluorescence analyses were carried out using a PerkinElmer LS 55 fluorescence spectroscopy and scan speed and excitation slit width were set at 500 and 5 nm, respectively. In intrinsic fluorescence after incubation of HSA with ZnO NPs, fluorescence intensity of tryptophan (Trp) residue was measured in the absence and presence of different concentrations of ZnO NPs. The excitation wavelength was adjusted at 295 nm and emission spectra were recorded in the wavelength range of 300-500 nm, and so the Stern-Volmer equation was used to determine the mechanism of quenching as following ^[11]^:

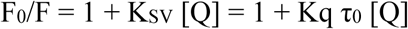

Where F_0_ and F are the fluorescence intensity of HSA in the absence and presence of quencher (ZnO NPs), respectively. K_SV_ is the Stern-Volmer quenching constant, [Q] is the concentration of quencher, Kq is the quenching rate constant and τ_0_ is the average lifetime of HSA in the absence of quencher and its value is 10^−8^ s ^[12]^. In addition to Trp emission, also ANS fluorescence probe was used for monitoring conformational changes in HSA ^[13]^. First, HSA was incubated with ANS (3 μM) and different concentrations of ZnO NPs, and then fluorescence intensity of ANS was measured in the absence and presence of different concentrations of ZnO NPs in the wavelength range of 400-600 nm, upon excitation at 350 nm.

### 2.4. Theoretical procedures

In order to investigate the interaction details of ZnO NP with HSA at the atomistic level, md simulations were carried out using the Amber99SB force field ^[14]^, in Gromacs package, version 2018 ^[15]^. The crystallographic structure of human serum albumin (PDB ID: 1e7e) was obtained from Protein Data Bank ^[16]^. To understand the mechanism of ZnO NP effect on apo-HSA structure, theoretical studies were done in absence and presence of ZnO NP. In the all md simulations was used from a spherical ZnO NP with diameter of 4 nm which has hexagonal wurtzite-type crystal structure with the lattice constants a=3.25 Å and c=5.2 Å ^[18]^. The interactions between Oxygen and Zinc atoms are described by Lennard-Jones potentials and the values of σ and ε parameters respectively are set to 2.128 Å and 0.418 kJ.mol^−1^ for the O atom and 1.711 Å and 1.254 kJ.mol^−1^ for the Zn atom. Partial charges (+1.026 and −1.026 for Zn and O atoms, respectively) were taken from the previous studies ^[19]^. In all simulations, protein and ZnO NP were placed in the center of a triclinic box with a distance of 1 nm from all edges, and under periodic boundary conditions (PBC). The distance between ZnO NP and HSA in the simulation systems was set to 6 Å. Then simulation boxes were filled by TIP3P water molecules ^[20]^ and the systems were neutralized by adding Na^+^ and Cl^−^ ions at the physiological concentration of 150 mM. The energy of the systems was minimized using the steepest descent algorithm and the minimization has stopped when the F_max_ was smaller than 10 kJ.mol^−1^.nm^−1^. Then, 100 ps NVT and 300 ps NPT ensembles were performed for each system until the temperature and pressure of the systems reach stable states in the 310 K and 1 atm, respectively. Also during the simulations was used from Particle Mesh Ewald (PME) method ^[21]^ and LINCS ^[22]^ and SETTLE ^[23]^ algorithms to calculate the long-range electrostatic interactions and constrain the length of bonds and position of the TIP3P water molecules, respectively. The cut-off for the coulomb and van der Waals interactions was set to 0.9 nm. Finally, all the systems in absence and presence of ZnO NP were simulated for 100 ns and with time step of 2 fs. The trajectories analysis was performed using GROMACS utilities and VMD and USCF Chimera ^[24]^.

## 3. Results and Discussion

### 3.1. Characterization of ZnO NPs

The SEM and TEM images of ZnO NPs are shown in (Figure 1a). The sheet like morphology of the ZnO NPs with approximate thickness less than 40 nm are seen.

**Figure 1.**
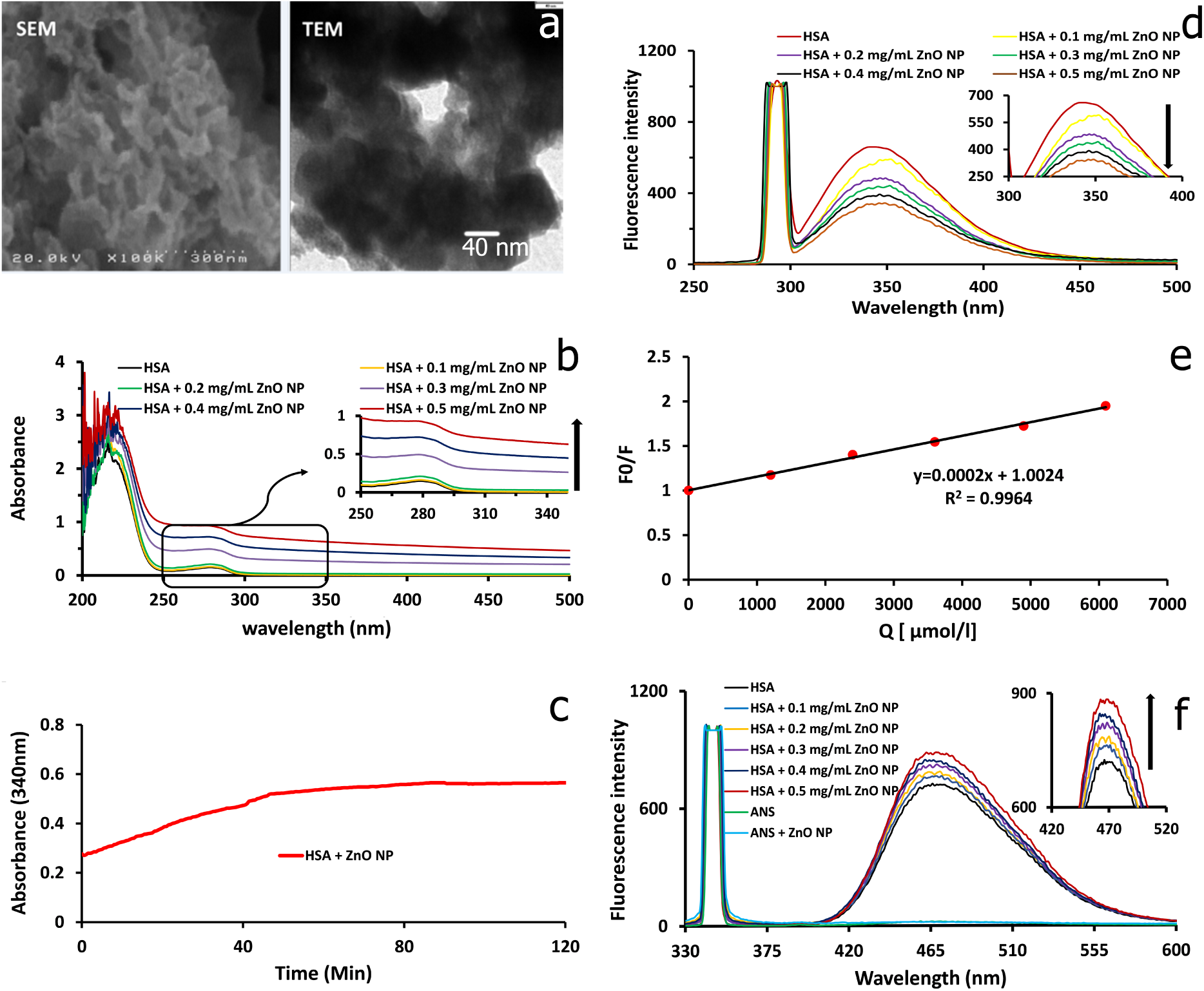
SEM and TEM micrographs of ZnO NPs (a). UV-Visible spectra of HSA in the presence of different concentrations of ZnO NPs (b). Time kinetics plot of HSA in the presence of 0.5 mg/mL of ZnO NPs (c). Intrinsic fluorescence spectra of HSA in the presence of increasing concentration of ZnO NPs. Inset shows enlarge view of the plot (d). The Stern-Volmer plot of fluorescence quenching of HSA by ZnO NPs (e). Extrinsic fluorescence spectra of HSA in absence and presence of different concentrations of ZnO NPs. Inset shows enlarge view of the plot (f).

### 3.2 UV-Vis spectroscopy and Time kinetics analyses

One of the most effective methods to study the structural changes in proteins is UV-Vis spectroscopy ^[25]^. UV-Vis absorption spectra of HSA in absence and presence of ZnO NPs are shown in (Figure 1b). The absorption spectra for HSA show two peaks at wavelengths of 230 and 280 nm. In order to investigate the conformational changes in the protein we studied the absorption spectra at 280 nm which corresponds to the π-π* electron transitions of aromatic amino acids (Trp, Tyr and Phe) ^[26]^. The absorption spectra of HSA at 280 nm gradually increased with the increasing of ZnO NPs concentration with no shift in absorption intensity. Similar results were observed in the articles which the ground state complex between proteins and NPs was formed ^[25, 27]^. Therefore, this increasing in the absorption intensity can be due to formation of the ground state complex between HSA and ZnO NPs. Also, it is possible that with the increasing of ZnO NPs concentration, it induced the structural changes in HSA and increased exposure of aromatic amino acid which followed with the increasing of adsorption intensity. Also, the time dependent absorption of HSA in the presence of ZnO NPs showed that the reaction between ZnO NPs and HSA, and consequently the conformational changes of HSA structure occur within the first 50 min, and thereafter no appreciable change in the absorption increasing occurs (Figure 1c).

### 3.3. Fluorimetric analyses

Intrinsic or natural fluorescence of proteins originates with the three aromatic amino acid residues include tryptophan, tyrosine and phenylalanine. HSA has only one tryptophan which present in the bottom of hydrophobic pocket in domain IIA (Trp 214). This residue is highly sensitive to its local micro-environment and a trivial change in the local environment either by ligand binding or conformational changes would significantly quench it ^[9, 25]^. Phenylalanine has a very low quantum yield and the fluorescence of tyrosine can be quenched if it’s ionized or placed near an amino group, a carboxyl group or a tryptophan ^[28, 29]^. For identifying variations in HSA folding, fluorescence emission spectra of tryptophan in absence and presence of ZnO NPs were measured using fluorescence spectroscopy (Figure 1d). The emission intensity of Trp decreases with an increase in ZnO NPs concentration which indicates quenching of the intrinsic HSA fluorescence by ZnO NPs. Previous works have reported similar quenching on the interaction of proteins with NPs ^[26, 29]^. According to the Stern-Volmer plot (Figure 1e), the plot of F0/F against [Q] is a straight line and slope of line is K_SV_. Therefore, was calculated a value of 2 × 10^2^ M^−1^ for K_SV_ and using the τ_0_ = 10^−8^ s was calculated 2 × 10^10^ M^−1^s^−1^ for Kq. To study the extrinsic fluorescence, we used from ANS that is a fluorescent probe and binds to hydrophobic pockets on the surface of proteins, that in HSA, it binds mainly to subdomain IIIA and formed ANS-HSA complex ^[13, 30]^. Therefore, the change in the ANS emission spectra gives useful information about conformational change of protein. ANS alone does not show any fluorescence emission, while the binding of it to HSA shows the fluorescence emission intensity gradually increased along the increasing of ZnO NPs concentration (Figure 1f). This indicated that with the increasing of ZnO NPs concentration, occurred structural changes in HSA and exposed hydrophobic pockets of protein that were buried in the interior of HSA molecule, and then the binding of ANS to these pockets along emission intensity increased.

### 3.4. Molecular dynamics simulation analyses

To investigation the stability and dynamics of HSA, we used Root Mean Square Deviation (RMSD) analysis, which show the conformational and orientational changes of the protein structure by comparing the displacement in the position of protein atoms relative to the reference structure ^[31]^. According to Figure 2a, RMSD values in presence of ZnO NP slightly increased that show the minor structural changes in protein during the 100 ns simulation.

**Figure 2.**
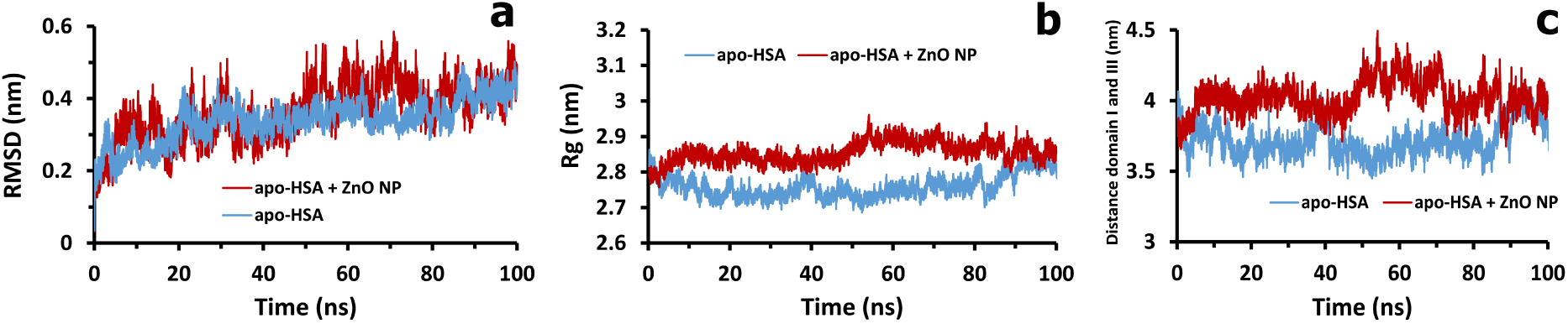
Backbone RMSD plot of apo-HSA in the absence and presence of ZnO NP (a). RG plot of HSA backbone (b) and distance domain I and III plot of HSA at different simulation systems during the 100 ns simulation (c).

Also, to measure the compactness of HSA structure we used Rg analysis that is defined as the approximate distance of atoms from the center of protein. The Rg value of HSA in presence ZnO NP is more, that is the reason for loss of protein compactness during the 100 ns simulation (Figure 2b). Also, distance domain I and III of HSA analyses show in presence ZnO NP increase the distance between domain I and III that is another reason for loss of protein compactness during the 100 ns (Figure 2c).

In addition, we investigated the adsorption of HSA on the surface of nanoparticle. The results showed that the distance between them decreased to less than 0.4 nm and remained constant along simulation time, which indicates electrostatic interactions between Zn and O atoms with acidic and basic residues, respectively. In Figure 3a is shown distance between one of the acidic and basic residues of HSA with Zn and O atoms, respectively. Therefore, electrostatic interactions are the most important factor in the adsorption of HSA on the surface of ZnO NP and the formation of nanoparticle-protein corona (Figure3b).

**Figure 3.**
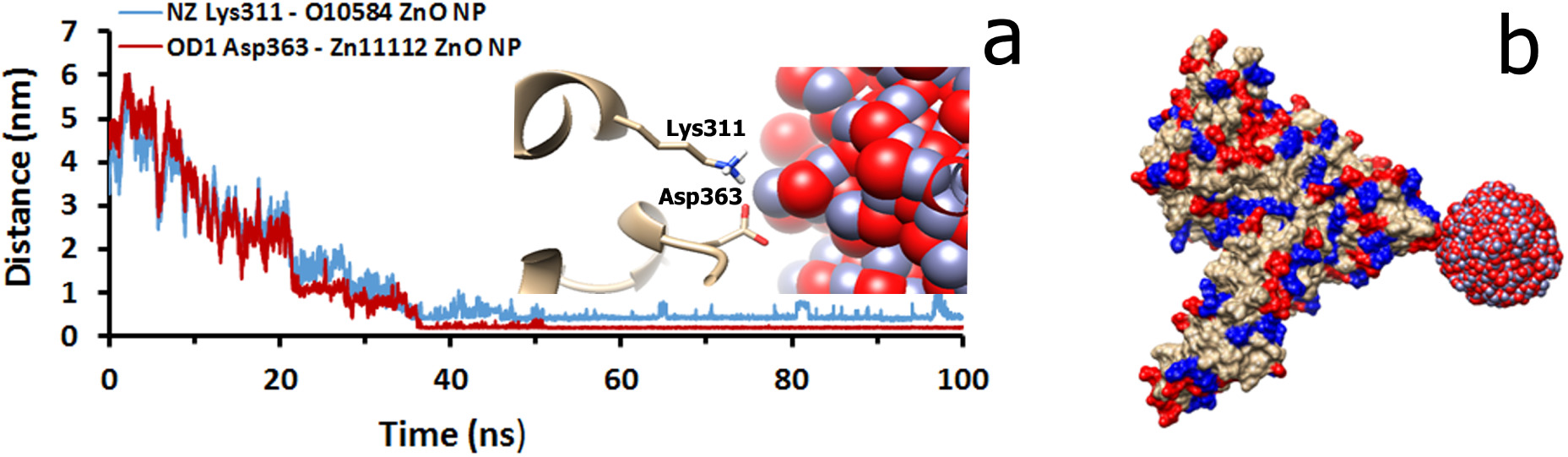
Distance between Lys311 and Asp363 of HSA with O and Zn atoms of ZnO NP, respectively, during the 100 ns simulation. Inset shows distance between them in last frame of simulation (O and Zn atoms are illustrated as red and blue colors, respectively) (a). The presentation of non-covalent interactions between HSA and ZnO NP and the formation of nanoparticle-protein corona. Basic and acidic residues of HSA are illustrated as blue and red colors in surface presentation, respectively, and other residues are shown in gray (b).

## 4. Conclusion

In the present study, the toxicity of ZnO NPs on HSA was investigated by *in vitro* and *in silico* methods. In *in vitro* studies, the ZnO NPs were synthesized with size less than 40 nm and the morphology of the nanosheet by hydrothermal method. The results of the UV-Vis and fluorescence spectroscopy showed the conformational changes of HSA in presence of ZnO NPs with the increasing of ZnO NPs concentration. Also, the Stern-Volmer plot showed that the intrinsic fluorescence of HSA was quenched through static mechanism. Furthermore, the results of md simulation indicated that ZnO NP causes minor structural changes in the protein structure during the 100 ns simulation, and the formation of nanoparticle-protein corona. Therefore, considering the numerous important functions of albumin in blood, the formation of nanoparticle-protein corona leads to loss the function of protein. Furthermore, with the passage of time, due to the abundance of acidic and basic residues in HSA structure −97 acidic and 83 basic amino acid- and also ZnO NP surface charges and the formation of high attractive and repulsive forces between them, unfolding of HSA is possible. So, it can be concluded ZnO NP lead to the structural and the functional changes of HSA and consequently, they may have the toxic effect on the health.

## Acknowledgement

This investigation was supported by a grant from Golestan University, Gorgan, Iran.

